# PRAME expression in melanoma is negatively regulated by TET2-mediated DNA hydroxymethylation

**DOI:** 10.1101/2024.07.26.605293

**Authors:** Rui Fang, Tuulia Vallius, Arianna Zhang, Devon Van Cura, Francisco Alicandri, Grant Fischer, Elizabeth Draper, Shuyun Xu, Roxanne Pelletier, Igor Katsyv, Peter K. Sorger, George F. Murphy, Christine G. Lian

## Abstract

Preferentially Expressed Antigen in Melanoma (PRAME) and Ten-Eleven Translocation (TET) dioxygenase-mediated 5-hydroxymethylcytosine (5hmC) are emerging melanoma biomarkers. We observed an inverse correlation between PRAME expression and 5hmC levels in benign nevi, melanoma in situ, primary invasive melanoma, and metastatic melanomas via immunohistochemistry and multiplex immunofluorescence: nevi exhibited high 5hmC and low PRAME, whereas melanomas showed the opposite pattern. Single-cell multiplex imaging of melanoma precursors revealed that diminished 5hmC coincides with PRAME upregulation in premalignant cells. Analysis of TCGA and GTEx databases confirmed a negative relationship between TET2 and PRAME mRNA expression in melanoma. Additionally, 5hmC levels were reduced at the PRAME 5’ promoter in melanoma compared to nevi, suggesting a role for 5hmC in PRAME transcription. Restoring 5hmC levels via TET2 overexpression notably reduced PRAME expression in melanoma cell lines. These findings establish a function of TET2-mediated DNA hydroxymethylation in regulating PRAME expression and demonstrate epigenetic reprogramming as pivotal in melanoma tumorigenesis.

**Teaser:** Melanoma biomarker PRAME expression is negatively regulated epigenetically by TET2-mediated DNA hydroxymethylation

## INTRODUCTION

Accurate and reproducible histopathologic diagnosis of melanoma, one of the most aggressive forms of skin cancer, is challenging and has profound clinical impact. Early and accurate diagnosis is crucial for effective management and improved patient outcomes. In this context, biomarkers play an important role but it remains unclear which biomarkers are most effective for diagnosis. Preferentially Expressed Antigen in Melanoma (PRAME) has recently emerged as a candidate diagnostic biomarker that can be detected using standard immunohistochemistry (1–4). Studies have shown immunostains for PRAME to represent a means of ancillary histopathology testing with high sensitivity and specificity for melanoma diagnosis, including ambiguous melanocytic neoplasms (1, 5, 6) and rare subtypes of melanoma (7–10). This specificity arises because PRAME, a membrane-bound antigen, is not typically expressed in normal cells except for the testis (11, 12). However, PRAME activation directly correlates with adverse clinical outcomes in several cancer types and is thought to contribute to tumorigenesis through inhibition of the retinoic-acid pathways for growth arrest and apoptosis (13, 14). Due to its differential expression profile and critical role in cancer progression (12, 13), PRAME has also garnered interest as a potential as a therapeutic target (15–20). However, despite its diagnostic and potential therapeutic importance (21–23), the biological mechanisms governing PRAME over expression in cancer have remained largely unknown since its original discovery in 1997 (11).

The TET family of Fe(II)/2-oxoglutarate dependent dioxygenases plays a critical role in mammalian gene regulation by catalyzing removal of epigenetic methylation marks from 5-methylcytosine (5mC) through successive oxidation reactions (24–26). The first product of this pathway, 5-hydroxymethylcytosine (5hmC) is a key intermediate for DNA demethylation and may also function as a stable epigenetic mark with context-dependent regulatory functions (27–29). Our prior research identified Ten-Eleven Translocation dioxygenase 2 (TET2) mediated loss of 5hmC marks as a functionally significant epigenetic hallmark of melanoma (30). Further studies have validated our findings, confirming that DNA hydroxymethylation plays an important role in melanoma tumorigenesis and that loss of 5hmC is a biomarker of melanocytic dysplasia, melanoma precursor lesions (31, 32), and malignant transformation in melanoma (33–36) and other cancers (29, 37, 38). Epigenetic reprogramming by the DNA methyltransferase inhibitor 5-aza-2’-deoxycytidine (5-azaC) reverses the global loss of 5hmC with prognostic implications in ovarian cancer (39).

Several studies suggest that DNA hypomethylation is associated with up-regulation of PRAME expression in acute myelogenous leukemia (AML) (40), MDS (41, 42), and ovarian cancer (43, 44). Other studies have shown that PRAME expression can be induced by 5-azaC treatment in AML (45) and melanoma (46). A recent study also shows that combining immunostains for PRAME and 5hmC increases sensitivity as compared to using PRAME alone for melanoma diagnosis (47). However, the potential role of 5hmC in regulating PRAME expression in melanoma remains unexplored. We hypothesized that inactivation of TET and the resulting loss of 5hmC marks might be trigger for PRAME overexpression during melanoma pathogenesis; in this study we test this idea. We demonstrate a significant inverse correlation between PRAME expression and 5hmC levels by using immunohistochemistry (IHC) and immunofluorescence (IF) in clinically annotated melanoma cohorts. TET2-overexpression (OE) in cultured human melanoma cell lines resulted in a significant decrease in PRAME expression and this reduction is associated with increased 5hmC levels. The effect of 5hmC on PRAME expression is likely direct: genome-wide mapping of 5hmC using hydroxymethylated DNA immunoprecipitation sequencing (hMeDIP-seq) reveals significantly increased 5hmC binding at the PRAME 5’ promoter region in TET2-OE cells as compared to control melanoma cells. Thus, PRAME expression in melanoma can be negatively controlled by TET2-mediated 5hmC epigenetic modification. This insight enhances our understanding of melanoma biology, potentially reveals new therapeutic targets, offering novel directions for further research and therapeutic strategies.

## RESULTS

### Reciprocal immunostaining pattern of PRAME and 5hmC in human melanocytic neoplasms

Multiplex IF for 5hmC (orange), PRAME (red), the melanocytic lineage marker MART-1 (green), and DAPI (blue) was performed on a human melanoma tissue microarray (TMA) that contains duplicated core tissue from benign nevi (n=7), melanoma in situ (MIS) (n=3), primary cutaneous melanoma (n=18) and metastatic melanoma (n=13) (48). We quantified PRAME expression as the percentage of positive cells among melanocytic cells that were also MART^+^ ; we then scored the intensity of 5hmC staining in a 0-12 scale, as previously described (30). We observed negative PRAME staining and positive 5hmC staining (PRAME^-^/5hmC^high^) in benign nevi, and the opposite pattern of low 5hmC and high PRAME (PRAME^+^/5hmC^low^) in MIS, primary and metastatic melanomas (**Figure 1A** and **supplementary Figure 1A**).

**Figure 1.**
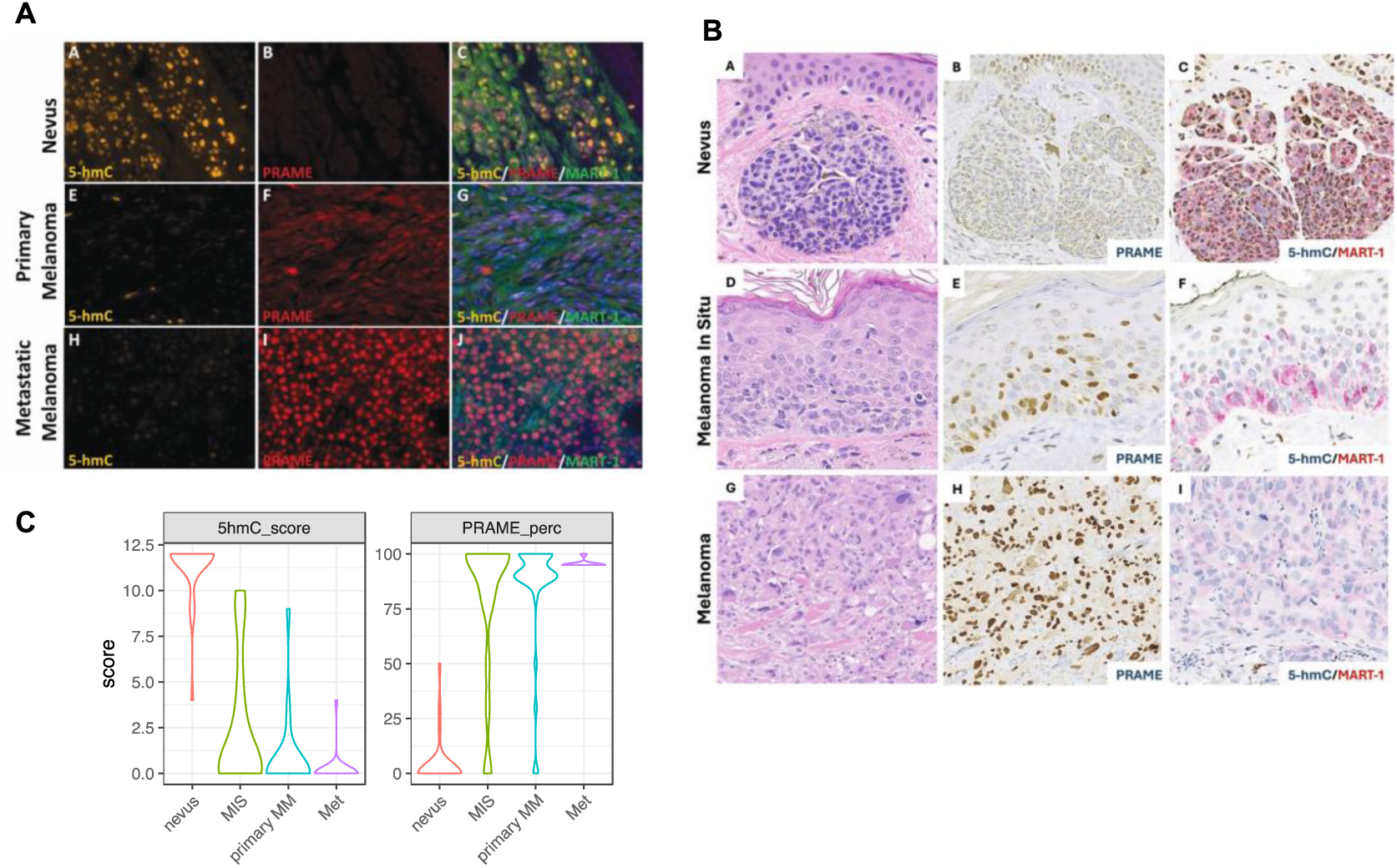
The inversed PRAME and 5-hmC staining in nevi and melanoma. **A.** Representative multiplex immunofluorescence staining of 5hmC (yellow), PRAME (red), MART-1 (green) in nevi, primary and metastasis melanoma, and counter staining with DAPI (blue). **B.** Representative IHC staining of nevus, melanoma in-situ and primary malignant melanoma. Panel a, d &g, H&E staining; panel b, e & h, PRAME, brown; panel c, f & I, dual IHC for 5hmC (nuclear positivity, brown) and melanocytic lineage marker MART-1 (membranous stain, red). **C.** Violin plot of PRAME and 5hmC scores combining cohort in A and B. PRAME is scored byt the percentage of PRAME positive cells. 5hmC, by either IF or IHC, are scored in a scale of 0-12 using a scheme based on intensity and percentage of positive cells. Nevus n=30; MIS n=19; Primary MM (primary malignant melanoma) n=38; Metastatic melanoma n=24.

To validate these findings, we examined a second cohort of whole slide melanocytic neoplasms stained by the BWH pathology core service using conventional one or two-color IHC; the cohort included nevi (n=20), MIS (n=23), primary cutaneous malignant melanoma (n=13) and metastatic melanomas (n=3). IHC for PRAME (brown nucleus and cytoplasm positivity) as well as dual IHC for 5hmC (brown nuclear positivity) and MART-1 (red membrane positivity) (**Figure 1B**) were performed on all specimens and evaluated using a 0-12 grading scale that was based on percentage of 5hmC positive cell counts and intensity (as previously described) (30, 33–35). PRAME IHC was reviewed by three pathologists, and the status of 5hmC staining was assessed by three additional researchers without knowledge of the clinical and pathologic features. Negative controls were run concurrently for all specimens and scored as < 1% nuclear staining. PRAME positivity was scored using a 75% cutoff as in literature (3, 5, 49, 50). Consistent with results obtained by IF, we observed a PRAME^-^/5hmC^+^ phenotype in benign nevi, and PRAME^+^/5hmC^low^ phenotype in the majority of melanoma cases (**Figure 1B**). However, two melanoma cases with nevoid morphology were PRAME^-^, consistent with the previously reported loss PRAME immunoreactivity in nevoid and desmoplastic melanoma (47, 51); 5hmC levels were high in our nevoid melanomas, further demonstrating the inverse correlation between PRAME and 5hmC (combined results of the two cohorts are summarized in **Figure 1C, supplementary Figure 1C** and **supplementary Table 1)**.

In addition, we observed a more diverse staining pattern in MIS than in more fully evolved primary and metastatic lesions. While the majority of MIS cases (12/19) had a PRAME^+^/5hmC^-^ phenotype similar to primary and metastatic melanomas, two MIS cases were PRAME negative with high 5hmC, while other MIS cases (5/19) exhibited variable PRAME positivity in combination with clearly detectable but reduced 5hmC staining.

### Correlation between PRAME expression and 5hmC levels in initiation and progression of human melanomas

The finding of variable PRAME and 5hmC levels in MIS prompted us to explore a potential correlation between PRAME and TET2/5hmC levels in the premalignant to early malignant transitional phases of melanoma evolution. A cohort of primary melanoma samples from 14 patients were examined using cyclic immunofluorescence (CyCIF) imaging and antibodies against SOX10 (melanocytes), CK14 and panCK (epidermal keratinocytes), PRAME, TET2 and 5-hmC (52–54). These patient samples were selected because they had contiguous fields of normal melanocytes bordering regions of melanocytic atypia that were directly adjacent to MIS and invasive areas. These progression-related histologic tissue domains (regions of normal, precursor, melanoma in situ, radial and vertical growth phase melanoma) were annotated using H&E-stained sections, and a total of 32 precursor regions were identified by histopathological re-review (**Figure 2A**). After single-cell segmentation and quality control, we identified 2,511 melanoma precursor cells positive for melanocytic lineage marker SOX10 (**Figure 2B**). In regions of melanocytic atypia, SOX10^+^ cells exhibited a negative correlation between PRAME and 5hmC levels at both a regional level (chi-square p=0.02) and in single cells (r=-0.24, p<0.001, **Figure 2C and 2D**). Based on these data we conclude that an inverse relationship exists between 5hmC and PRAME expression. This ranges from a PRAME negative and 5hmC high state in normal melanocytes and non-cancerous nevi to loss of 5hmC expression and induction of PRAME activation in overt melanomas; melanoma precursor cells exhibit an intermediate phenotype with partial loss of 5hmC expression and increasing levels of PRAME. Thus, the switch from a PRAME^-^/5hmC^+^ to a PRAME^+^/5hmC^-^ phenotypes appears to be graded and correlate with increasing melanoma stage.

**Figure 2.**
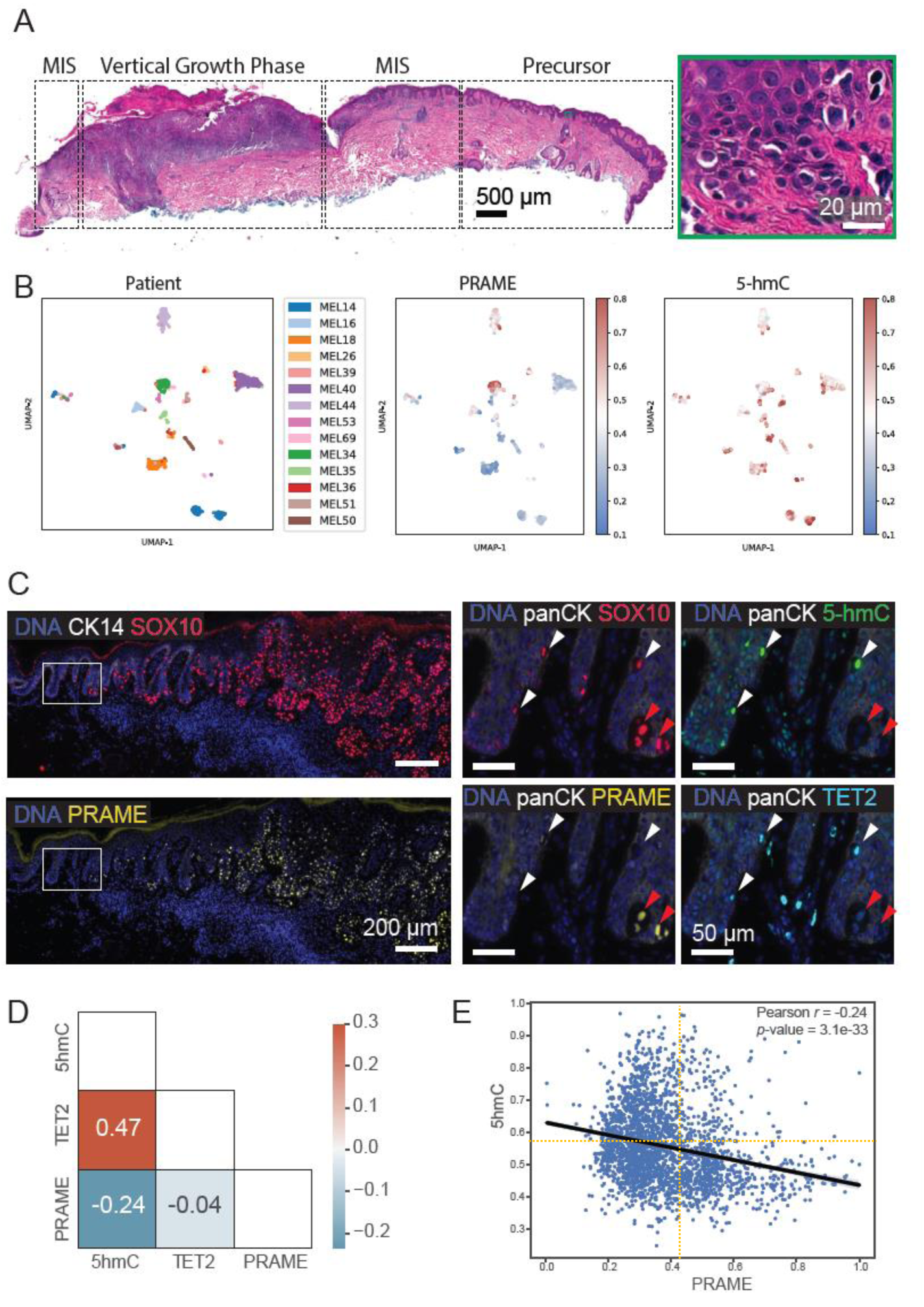
Inverse correlation of PRAME and 5hmC in melanoma precursors. **A.** An exemplary case (MEL18) highlighting the presence of progression-related histopathologic tissue regions (precursor, MIS) adjacent to a region of vertical growth phase melanoma. A proportion of the region with melanocytic atypia is magnified (green inset). Scale bars, 500 μm and 20 μm. **B.** Uniform manifold approximation and projection (UMAP) of single-cell data derived from CyCIF of 14 patient samples, labeled by patient ID (left) and the signal intensities of PRAME (middle) and 5hmC (right). **C.** CyCIF image of sample MEL14 showing the transition from precursor to melanoma in situ and invasion stained for DNA (blue), CK14 (white), SOX10 (red, top left) and PRAME (yellow, bottom left). The inset squares correspond to magnified panels on the right, with DNA (blue), PanCK (white), SOX10 (red), PRAME (yellow), 5hmC (green) and TET2 (cyan), highlighting the PRAME+ and PRAME-atypical melanocytes in the precursor region. Scale bars, 200 μm and 50 μm. **D.** Pearson correlation of PRAME, TET2 and 5hmC at the single cell level. **E.** Scatter plot of PRAME and 5hmC CyCIF mean fluorescence intensities of 2,511 SOX10^+^ single melanoma precursor cells in 32 precursor regions from 14 patients.

### Negative regulation of PRAME gene expression via TET-mediated DNA hydroxy-methylation and 5’ promoter region binding in melanocytic neoplasms

It is known that both PRAME activation and 5hmC loss play important roles in driving melanoma development (1–4, 7–10, 30–36). The inverse relationship between these markers in both melanoma precursor fields and during melanoma evolution suggests that 5hmC may regulate PRAME gene expression directly. To investigate this, we examined hMeDIP-Seq data for both nevi and melanomas (30). We observed a marked reduction in 5hmC binding to the 5’ promoter region of the PRAME gene in melanomas compared to nevi (**Figure 3A**). 5hmC reduction was also observed in the gene body and 3’ downstream regions of PRAME. Using independent nevi and melanoma samples, we also performed hMeDIP-qPCR and confirmed significant 5hmC reduction at the CpG island and the PRAME promoter (**Figure 3B**).

**Figure 3.**
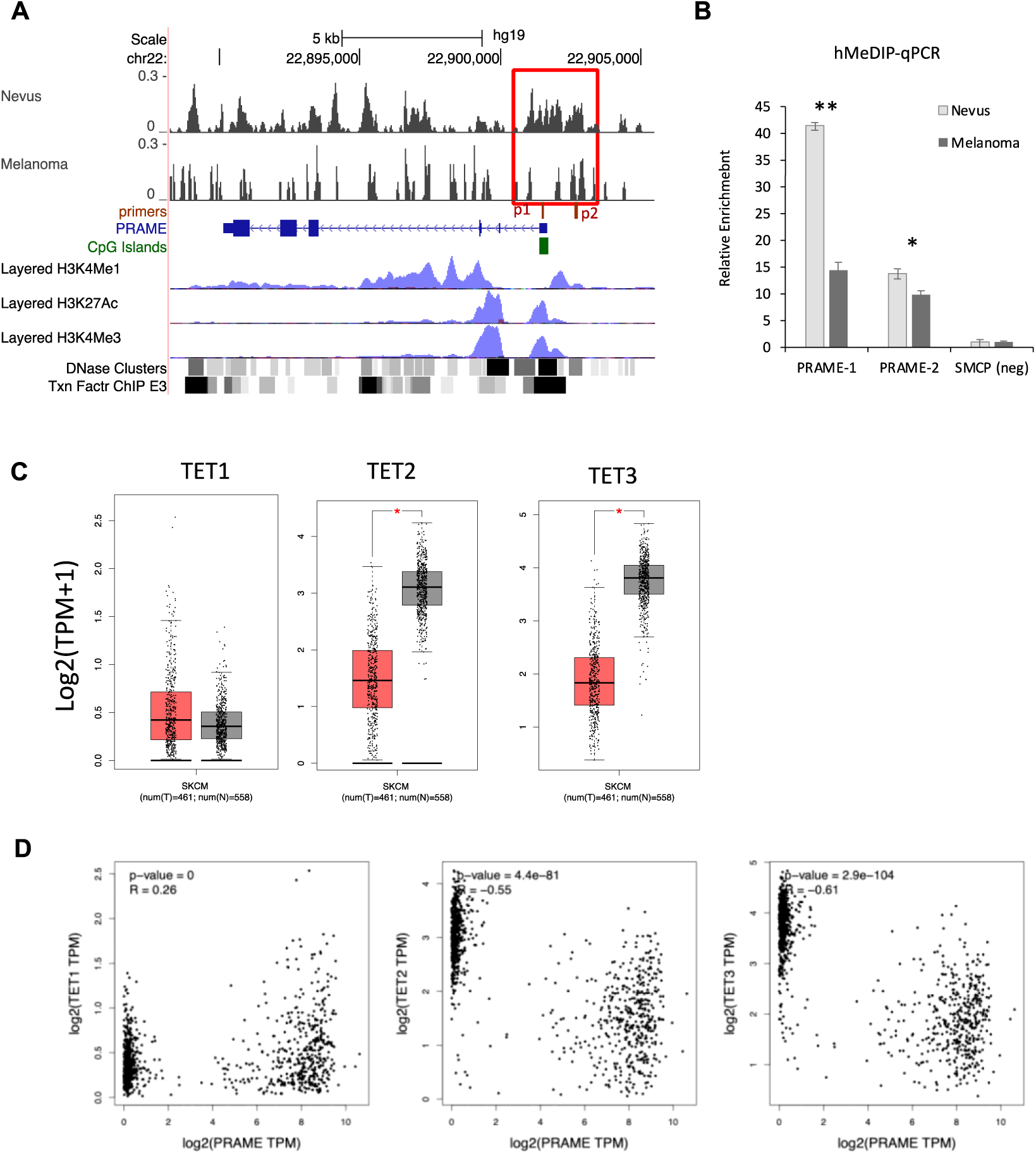
5hmC levels are reduced the PRAME promoter, concurring with the down regulation of TET2 and TET3 gene expression in melanoma. **A.** hMeDIP-seq detected decreased 5hmC levels at the 5’ promoter region of the PRAME gene in melanoma compared to nevi samples. Boxed region, 5hmC sites at PRAME promoter shown 5hmC loss in melanoma. Red bars, position of hMeDIP-qPCR primers in (B). The layered H3K4me1, H3K4me3 and H3K27ac from The Encyclopedia of DNA Elements (ENCODE) mark promoter and potential enhancers. **B.** hMeDIP-qPCR validation using independent melanoma and nevi samples. **, p<0.001; *, p<0.05. Error bar, standard deviation of triplicate qPCR reactions. Enrichment relative to SMCP 5hmC negative control region and normalize by inputs. **C.** Boxplot of TET1/2/3 expression in healthy and melanoma samples in the TCGA and GTEx RNA-seq databases. *, p<0.001 **D.** Correlation plot of PRAME with TET1, TET2, and TET3 gene expression in healthy and melanoma in the TCGA and GTEx RNA-seq databases. R, Pearson correlation coefficient.

5hmC is the oxidation product of 5mC and the conversion is mediated by the Ten-Eleven Translocation dioxygenases, TET1, TET2 and TET3. TET2 is a tumor suppressor gene frequently mutated or down-regulated in melanoma, AML and many other types of cancers (30, 38, 55). Examining RNA-seq datasets from The Cancer Genome Atlas (TCGA) and Genotype-Tissue Expression (GTEx) databases, we observed significant down regulation of TET2 and TET3 in melanomas (n=460) as compared to healthy skin samples (n=551) (**Figure 3C**). In contrast, the inferred level of expression of TET1 was lower than TET2 and TET3 and did not change between melanoma and normal skin, suggesting a less important role in regulating global 5hmC levels. Moreover, PRAME gene expression negatively correlated with TET2 (Pearson coefficient r= -0.55, p=4.4×10^-81^) and TET3 levels (r= -0.61, p=2.9×10^-104^), consistent with the observed inverse correlation between 5hmC and PRAME measured by IF and IHC (**Figure 3D)**; TET1 positively correlated with PRAME levels in the same analysis; r=0.26, p=0) (**Figure 3D**). We therefore hypothesized that TET2 and/or TET3 regulates PRAME gene expression by functioning as transcriptional suppressors.

### Restoration of 5hmC levels and decreased PRAME expression by TET2-overexpression in cultured human melanoma cells *in vitro*

To test this hypothesis, we expressed TET2 ectopically in melanoma cell lines. We have previously reported that TET2 overexpression restores the 5hmC landscape along the genome of A2058 human melanoma cells and reduces tumor growth *in vivo* (30). We therefore over-expressed full-length TET2 (TET2-OE) in A2058 and performed hMeDIP-seq (**Figure 4A**) and 5hmC DIP qPCR (**Figure 4B**) to measure 5hmC levels at the PRAME gene promoter. We found that 5hmC levels at the PRAME promoter were significantly higher in TET2-OE cells.

**Figure 4.**
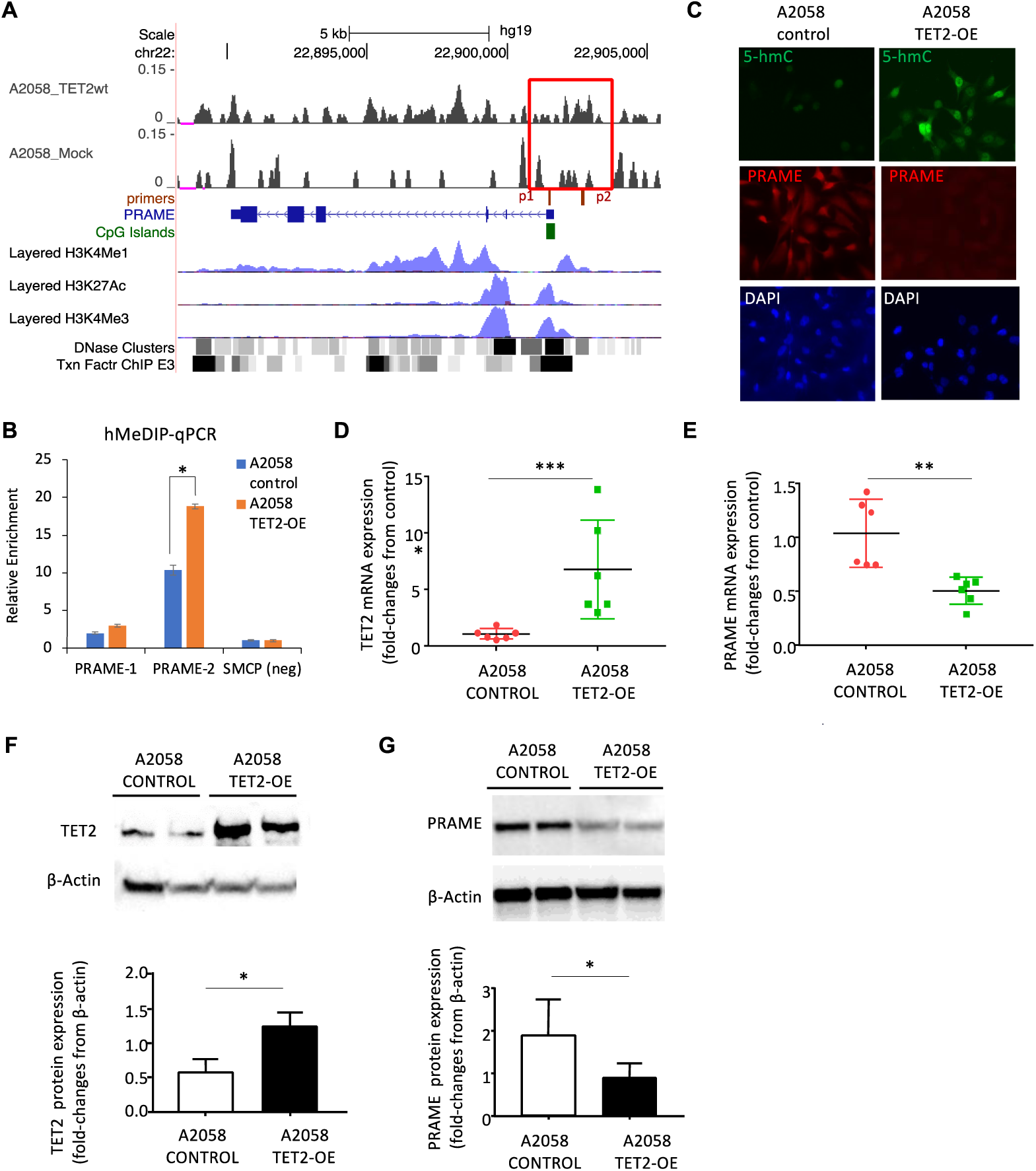
TET2 ectopic expression restores 5hmC at PRAME promoter and suppresses PRAME gene expression. **A.** hMeDIP-seq profile at PRAME genes in A2058 control and A2058 TET2-OE cells as in Fig. 3A. **B.** hMeDIP-qPCR validation using independent samples. *, p<0.05. Error bar, standard deviation of triplicate qPCR reactions. Enrichment relative to SMCP 5hmC negative control region and normalize by inputs. **C.** Immunofluorescence staining of 5hmC (green) and PRAME (red) A2058-Control and A2058-TETOE counter staining with DAPI (blue) **D-E.** Comparison of TET2 (B) and PRAME (C) mRNA levels in A2058-Control cells and A2058-TET2OE cells by qRT-PCR. **F-G.** Western blot of TET2 (F) and PRAME (G) in A2058-Control and A2058-TET2OE cell lines. β-actin was utilized as a loading control. Bottom panel, densitometry quantification of western blots. *, p<0.05; **, p<0.01; ***, p<0.001, by Student’s T-test.

We next performed IF staining and confirmed significantly higher global 5hmC levels in A2058-TET2OE cells as compared to A2058 control cells (**Figure 4C**). A2058-TET2OE cells also stained strongly for PRAME but PRAME was undetectable in parental A2058 cells (**Figure 4C**). RT-qPCR analysis confirmed these findings by showing that A2058-TET2OE cell line had significantly higher expression of TET2 (p<0.001, **Figure 3D**) and significantly lower expression of PRAME mRNAs as compared to A2058-control cells (p<0.01, **Figure 4E**).

Western blot analysis served as a final confirmation: TET2 protein levels were significantly higher and PRAME levels lower in A2058-TET2OE as compared to parental cells (p<0.05, **Figure 4F**, **4G**). Taken together, these results demonstrate that restoration of 5hmC marks by ectopic TET2 expression in cultured melanoma cells suppresses PRAME gene and protein expression levels.

## DISCUSSION

Using pathophysiologic studies of clinical samples and molecular studies in cell lines, we have established a novel and potentially important function forTET2-mediated DNA hydroxymethylation in negatively regulating PRAME expression in melanoma. These data suggest a cooperative role for 5-hmC loss and PRAME activation in driving melanoma transformation and progression. TET2 is significantly down-regulated in melanoma as compared to melanocytes in healthy skin, 5-hmC levels are lower, and PRAME expression is higher. This inverse correlation between high TET2 and 5-hmC on the one hand, and low PRAME expression on the other, is consistently observed at a whole-specimen level in melanoma clinical cohorts by IF and IHC for 5hmC, and by RNA-seq for TET2 (based on data in the TCGA and GTEx databases). Moreover, single-cell multiplexed image analysis using CyCIF shows that the inverse correlation between PRAME and 5hmC occurs as early as melanoma precursor cells in premalignant lesions. A direct connection between these changes is strongly suggested by our finding that ectopic expression of TET2 restores 5hmC marks at the PRAME promoter and down-regulates PRAME expression in cultured mammalian cells. Taken together, our data uggest that 5hmC loss, likely due to the down-regulation of TET2, plays an important role driving PRAME activation and that this may help drive melanoma malignant transformation and progression.

A recent paper by Kaczorowski and colleagues has reported loss of 5hmC and a concomitant increase PRAME expression in melanoma (56), which is consistent with our findings. However, the conclusion in Kaczorowski *et al* that PRAME expression is not correlated with DNA demethylase expression appears to be incorrect, and likely reflects that fact that the authors only stained for TET1 (by IHC) without assessing TET2 or TET3 expression. However, data in Figure 3D and analysis of RNA-seq data in the TCGA and GTEx databases show that TET2 and TET3, not TET1, correlate with PRAME gene expression. 5hmC and different members of the TET enzyme family may function differently because they promote DNA demethylation at different sites on the genome (57). Consistent with this idea, 5hmC is a stable marker in many cell types and specific TET enzymes are known to interact with numerous chromatin modifying factors and transcriptional co-activators and co-suppressors. These include the COMPASS complex and the NuRD and SIN3A/HDACs co-repressor complexes (58). Thus, 5hmC occupies a unique position in the genome, mediating gene activation and suppression depending on context. In melanoma, our data show that it is most likely TET2-mediated 5hmC production that suppresses PRAME gene expression.

In terms of clinical application of biomarkers, our findings align well with prior studies, particularly regarding the possibility of improved sensitivity and specificity of melanoma diagnosis using the simultaneous detection of 5hmC and PRAME (47). A notable aspect of our results are their implications for the diagnostic application of PRAME immunostaining, which has previously been limited to quantifying the number of cells that are positive or negative for PRAME for diagnostic purpose (3). In contrast, 5hmC immunostaining demonstrates a gradual decrease in staining intensity across the spectrum of melanocytic neoplasms, ranging from benign lesions to malignant melanomas (30, 33, 34, 36). This gradation in intensity offers a more detailed view on disease progression. Our methodology combines both the qualitative assessment of staining intensity and the quantitative measurement of cell positivity (30) and is a more comprehensive evaluation framework for melanoma diagnoses with enhanced accuracy and reliability (47, 51).

This study is limited by a relatively small sample size and a cohort derived from a single tertiary care institution. Future studies should consider including a larger set of cases from diverse populations and institutions. Rawson and colleagues demonstrated that the dual staining of PRAME and 5hmC increases diagnosis accuracy (47). However, limited by a relatively small sample sizes of our cohorts, we don’t have the statistical power to evaluate the benefit of PRAME/5hmC dual detection in melanoma diagnosis as compared to assessment of PRAME expression alone. Nevertheless, we have observed that most melanoma cases had significant loss of 5hmC and PRAME positivity, consistent with the reports in the literature (Supplementary Table 3). In addition, studies have shown that the frequency of PRAME positivity is not uniform across melanoma subtypes. For example, PRAME positivity is only present in approximately 35% of desmoplastic melanomas (3, 59); we observed that 5hmC levels are largely retained in desmoplastic melanoma (data not shown). Thus, further investigation of these two biomarkers is warranted in various subtypes of melanoma.

To our knowledge, ours is the first study to establish a link between aberrant PRAME expression in melanoma and underlying transcriptional regulators. These findings provide insight into melanoma epigenetics and underscore the diagnostic and prognostic utility of combined PRAME/5hmC staining. Future research should also focus on exploring the therapeutic potential of modulating TET2 activity and its impact on PRAME expression, potentially leading to novel strategies for blocking melanoma initiation and progression.

## MATERIALS AND METHODS

### Histopathologic Samples and Data

This study was conducted with approval of the Institutional Review Board of Massachusetts General Brigham, Harvard Medical School. An established tissue microarray was previously constructed from the archives of the Department of Pathology at Brigham and Women’s Hospital used and composed of a cohort of 41 cases of melanocytic lesions including benign nevi (n=7), melanoma in situ (n=3), primary cutaneous melanoma (n=18) and metastatic melanomas (n=13). Another cohort of 59 cases of melanocytic lesions, which included benign nevi (n=20), melanoma-in-situ (n=23), primary cutaneous melanoma (n=13) and metastatic melanomas (n=3) was obtained (Table 1). Approval for the archived cases and the tissue microarray study were granted by the Massachusetts General Brigham Human Research Committee. Informed consent was not necessary, as all tissue samples were deidentified. The hematoxylin and eosin stained sections and diagnosis were independently reviewed and confirmed by board-certified dermatopathologists (C.G.L, G.F.M, I.K.). We used Gene Expression Profiling Interactive Analysis (GEPIA) (60, 61) to examine the relationship between PRAME and the TET family of enzymes in public data repositories by analyzing the molecular profiles of melanoma and healthy skin samples in the Cancer Genome Atlas (TCGA) and the Genotype-Tissue Expression (GTEx) GTex database.

**Table 1.** Case cohort demographics.

|  | **Number of cases** | **Median age at diagnosis** | **Range of age at diagnosis, years** | **Sex, n (%)** | **Anatomic site, n** |
| --- | --- | --- | --- | --- | --- |
| **Nevus** | 21 | 57 | [26 - 79] | Male: 8 (38.1)<br>Female: 13 (61.9) | head and neck: 1<br>extremities: 5<br>trunk: 15 |
| **MIS** | 23 | 70 | [38 - 91] | Male: 9 (39.1)<br>Female: 14 (60.9) | head and neck: 6<br>extremities: 12<br>trunk: 5 |
| **Primary Melanoma** | 13 | 58 | [23 - 79] | Male: 7 (53.8)<br>Female: 5 (38.1)<br>Not reported: 1 (7.7) | head and neck: 2<br>extremities: 6<br>trunk: 4<br>not reported: 1 |
| **Metastatic Melanoma** | 3 | 63 | [44 - 83] | Male: 3 (100.0)<br>Female: 0 (0.0) | head and neck: 0<br>extremities: 1<br>trunk: 2 |

### Cell Culture

Human melanoma cell lines A2058 were obtained from ATCC (Manassas, VA). Melanoma stable cell lines overexpressing flag-tagged, full-length wild-type TET2 (TET2-OE) were established as described (30, 62). Briefly, A2058 control cells were transfected with empty vector plasmids. For the restoration of 5hmC levels, A2058 melanoma cells were transfected with plasmids encoding TET2 constructs using a standard lipofection technique (63). The overexpression of full-length TET2 and TET2-M control was verified by Western blot and quantitative RT-PCR, according to a standard protocol described by Lian et al (30, 62). All cells were grown in Dulbecco’s modified Eagle’s medium (Lonza, Hopkinton, MA). Culture media were supplemented with 10% heat-inactivated fetal bovine serum (HyClone, Logan, UT), 200 mmol/L L-glutamine, 100 IU/mL penicillin, and 100 mg/mL streptomycin (Life Technologies, Carlsbad, CA) and maintained at 37℃, 5% CO2.

### Western Blot

Western Blot was performed as previously described (48, 62, 64). Briefly, cell lysate was collected in 100μl lysis buffer (100mM Tris PH 6.8, 2% SDS and 12% glycerinum) and protein concentration was quantified using the Pierce BCA Protein Assay kit (Thermo Scientific, Catalogue#23227, Waltham, MA) 20 µg proteins were electrophoresed in running buffer on Mini-protean® TGXTM Precast Gel (Bio-Rad, Catalogue#456–1094, Hercules, CA) and transferred to Nitrocellulose membranes using transfer buffer. Primary antibodies—PRAME (Abcam, Catalogue#ab219650, Waltham, MA), TET2 (Proteintech, Catalogue#21207-1-AP, Rosemont, IL) were immunoblotted at 4 °C overnight on a shaker. After washing, the membranes were incubated with goat anti-rabbit IgG-HRP antibodies (Santa Cruz, Catalogue#sc-2357, Dallas, TX) at room temperature for 1 hour. Imaging and densitometry measurement were completed by ChemiDOCTM XRS+ with the Image Lab system (Bio-Rad, Hercules, CA).

### Quantitative Real-Time Polymerase Chain Reaction (RT-PCR)

Conditions for RNA extraction and quantitative RT-PCR analysis were implemented as previously documented (48). Briefly, total RNA of melanoma cells was extracted using the RNAeasy kit (Qiagen, Germantown, MD), and the cDNAs were synthesized using the SuperScript III First-Strand kit (Bio-Rad, Hercules, CA). Relative gene expression was normalized to β-actin). All reactions were run in triplicates Data was analyzed using the 2^−ΔΔCt (65). Primers used are reported in supplementary Table 2.

### Hydroxymethylated DNA Immunoprecipitation

Hydroxymethylated DNA immunoprecipitation sequencing (hMeDIP-seq) experiment was performed on human nevi, melanomas, and cultured human melanoma cell lines as previously described by Lian et al (30). Briefly, genomic DNA was purified from surgical tissues using Qiagen DNeasy Blood and Tissue kit and was sonicated to ∼200bp fragments using Covaris M220 Focused-ultrasonicator. dsDNA fragments (2μg) were denatured by heating to 98C for 10min and chilled on ice for 10min and incubated overnight with anti-5hmC antibody (4μl, Active Motif #39791) in 20mM Na-Phosphate pH 7.0, 0.14M NaCl, 0.05% Triton X-100. DNA-antibody complexes were captured using Dynabeads Protein A (Thermo Scientific) and enriched DNA was eluted and purified using Qiagen MiniElute PCR purification kit. hMeDIP-qPCR was performed using following primers specific to the PRAME promoter region (Figure 3B) and SMCP negative control which had low 5hmC. PRAME-p1 forward, 5’-CACTCTCCACAGAAATCCAC-3’; PRAME-p1 reverse, 5’-GCCCTGAATGTAGGGAAAG-3’; PRAME-p2 forward, 5’-GAAGGCAGCAGATGGATAG-3’; PRAME-p2 reverse, 5’-CAATGGAGCAACAGCAATC-3’; SMCP forward, 5’-GCTGAGTGGTTGAGTTTGAG-3’, SMCP reverse, 5’-GGGATTGTGCAGTTCTCTTC-3’. To quantify 5hmC changes, enrichment relative to SMCP 5hmC-negative control region were calculated using 2^-ΔCt^ and was normalized by corresponding input.

### Immunohistochemistry and Immunofluorescence

Immunohistochemistry (IHC) and Immunofluorescence (IF) were performed on 5 µm thick paraffin-embedded sections. Sections were treated with heat-induced antigen retrieval using target retrieval solution (Dako, Carpinteria, CA) and heated in a Pascal-pressurized heating chamber (Dako; 121℃ for 30 seconds, 90℃ for 10 seconds). After incubation at 4℃ overnight either with human PRAME antibody (Thermo Scientific, Catalogue#MA5-31408) or rabbit anti-5hmC (Active Motif, Carlsbad, CA), sections were incubated with horseradish peroxidase conjugated secondary antibodies for 30 minutes at room temperature, and signals were visualized with NovaRED horseradish peroxidase substrate (Vector Laboratories, Burlingame, CA), followed by a hematoxylin counterstain. In addition, triple color labeling immunofluorescence was performed to complement IHC analysis as a means of three-channel identification of epitopes, with cytoplasm staining for MART-1, nuclear staining for PRAME and 5hmC. Instead of incubation with the secondary antibodies, these sections were incubated with a mixture of goat anti-rabbit IgG (Thermo Scientific, Alexa Fluor 647-red) and goat anti-mouse IgG2b (Thermo Scientific, Alexa Fluor 488-green). Appropriate isotype matched antibody controls and tissue controls were used for all experiments. All specimens were evaluated separately according to a scoring system based on the percentage of positive cell counts. Intensity of staining of 5hmC IHC was determined on a semiquantitative scale of 0 to 3: no staining (score 0), weakly positive staining (score 1), moderately positive staining (score 2), and strongly positive staining (score 3). The entire area of two replicate 2-mm tissue microarray cores (6.28 mm2) was assessed. An immunoreactive score was derived by multiplying the percentage of positive cells by the staining intensity as previously described in literature (23–29).

### Cyclic immunofluorescence

Tissue-based Cyclic Immunofluorescence (CyCIF) was performed on 5 µm thick paraffin-embedded sections as previously described (52). In brief, the BOND RX Automated IHC Stainer was used to perform slide dewaxing and antigen retrieval. Slides underwent multiple cycles of antibody incubation, imaging, and fluorophore inactivation. Slides were imaged using a CyteFinder slide scanning fluorescence microscope (RareCyte Inc.) with a 20×/0.75 NA objective with no pixel binning. The list of the antibodies used in the CyCIF experiment is presented in Supplementary Table 3. Stitching individual images together into a high-dimensional representation for further single-cell segmentation and analyses was done using MCMICRO pipeline (54), an open-source multiple-choice microscopy pipeline (*version: 38182748aa0ec021f684ce47248c57340d2f4cc7 - full codes available on at* https://github.com/labsyspharm/mcmicro). Specific parameters were optimized after iterative inspection of results, to ensure accurate single cell identification (params.yml file). After generating the segmentation masks, the mean fluorescence intensities of each marker for each cell were computed, resulting in a single-cell data table for all whole-slide CyCIF images. The X/Y coordinates of annotated histologic regions on the whole-slide image were used to extract the single-cell data of cells that lie within the annotated precursor regions. Multiple approaches were also taken to ensure the quality of the single-cell data. At the image level, the cross-cycle image registration and tissue integrity were reviewed; regions that were poorly registered or contained severely deformed tissues and artifacts were identified, and cells within those regions were excluded. Antibodies that gave low confidence staining patterns by visual evaluation were also excluded from the analyses. After this, the single-cell phenotyping was performed as described (53), to identify melanocytes, and score the positivity of PRAME and 5hmC at both the regional and the single-cell level.

### Statistical Analysis

Statistical analyses were performed using appropriate software. The correlation between PRAME expression and 5hmC levels was assessed using Chi-Square and Pearson’s correlation coefficient. Differences in gene expression and 5hmC binding patterns between groups were evaluated using Student’s t-test or ANOVA, as applicable. A p-value of <0.05 was considered statistically significant.

## SUPPLEMENTARY MATERIALS

**Supplementary Table 1.** Summary of IF and IHC staining of nevus and melanoma patient samples.

| Staining | Diagnosis | PRAME% | 5hmC_score | differential PRAME alone (a,c) | differential dual biomarkers (a,b,c) |
| --- | --- | --- | --- | --- | --- |
|  | **27 nevus** | **0** | **12** | **PRAME-, benign** | **PRAME- AND 5hmC+, benign** |
| IHC | nevus | 50 | 4 | PRAME-, benign | PRAME-/5hmC-, melanoma |
| IHC | nevus | 30 | 12 | PRAME-, benign | PRAME- AND 5hmC+, benign |
| IHC | nevus | 20 | 9 | PRAME-, benign | PRAME- AND 5hmC+, benign |
|  | **12 MIS** | **>= 90%** | **0** | **PRAME+, melanoma** | **PRAME+ OR 5hmC-, melanoma** |
| IF | MIS | 95 | 6 | PRAME+, melanoma | PRAME+ OR 5hmC-, melanoma |
| IHC | MIS | 80 | 4 | PRAME+, melanoma | PRAME+ OR 5hmC-, melanoma |
| IHC | MIS | 80 | 3 | PRAME+, melanoma | PRAME+ OR 5hmC-, melanoma |
| IF | MIS | 50 | 10 | PRAME-, benign | PRAME-/5hmC+ benign |
| IHC | MIS | 30 | 2 | PRAME-, benign | PRAME+ OR 5hmC-, melanoma |
| IHC | MIS | 0 | 9 | PRAME-, benign | PRAME-/5hmC+ benign |
| IHC | MIS | 0 | 9 | PRAME-, benign | PRAME-/5hmC+ benign |
|  | **33 primary MM** | **>= 90%** | **0** | **PRAME+, melanoma** | **PRAME+ OR 5hmC-, melanoma** |
| IF | primary MM | 80 | 4 | PRAME+, melanoma | PRAME+ OR 5hmC-, melanoma |
| IHC | primary MM | 50 | 1 | PRAME-, benign | PRAME+ OR 5hmC-, melanoma |
| IHC | primary MM | 30 | 3 | PRAME-, benign | PRAME+ OR 5hmC-, melanoma |
| IHC | primary MM | 1 | 9 | PRAME-, benign | PRAME-/5hmC+ benign |
| IHC | primary MM | 0 | 9 | PRAME-, benign | PRAME-/5hmC+ benign |
|  | **23 Met** | **>= 90%** | **0** | **PRAME+, melanoma** | **PRAME+ OR 5hmC-, melanoma** |
| IF | MET | 95 | 4 | PRAME+, melanoma | PRAME+ OR 5hmC-, melanoma |

(a) PRAME<sup>+</sup>, >=75% PRAME positive cells
(b) 5hmC<sup>+</sup>, score >=4
(c) highlight, staining differential inconsistent with diagnosis.

**Supplementary Table 2.**
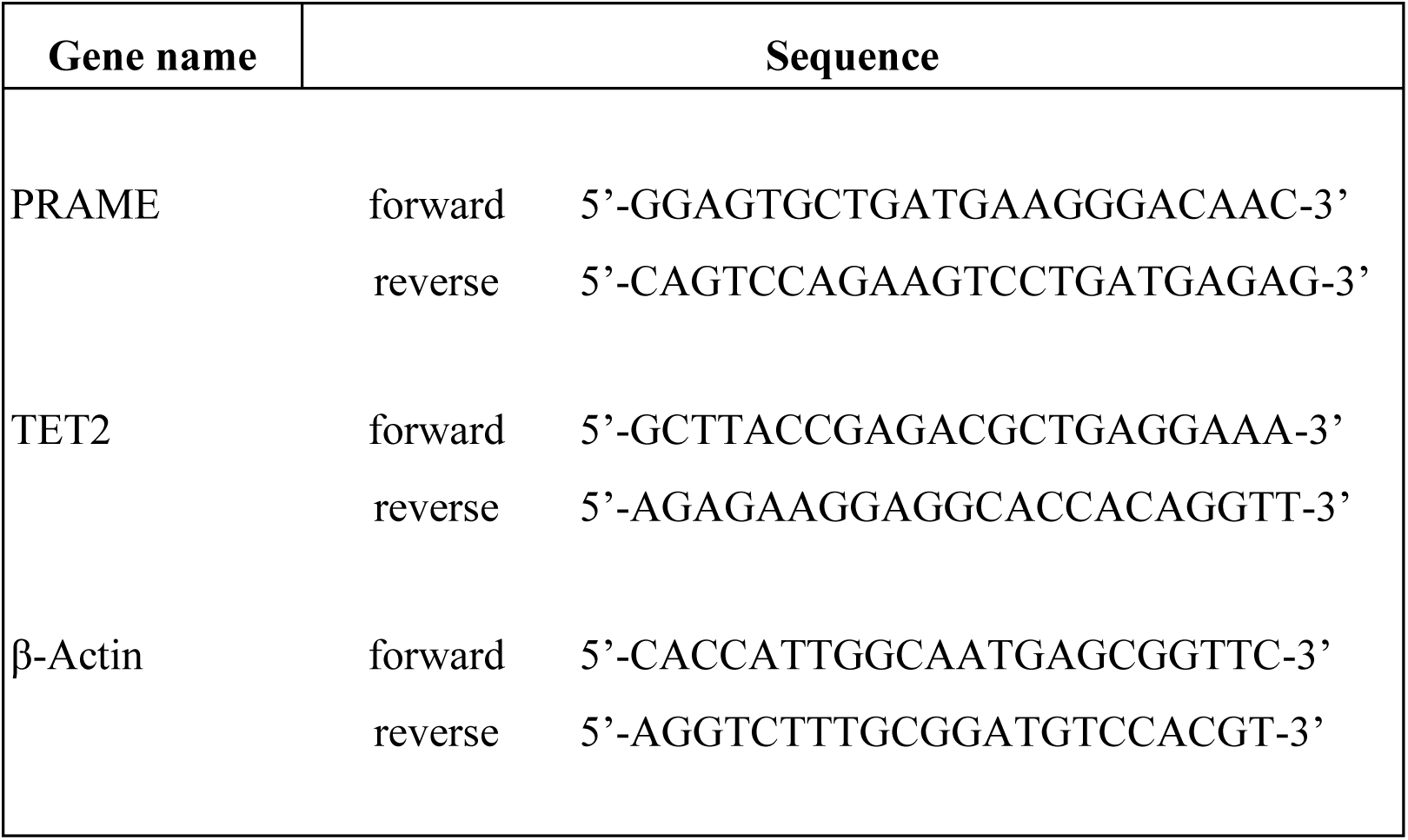
Gene-specific RT-qPCR primers.

**Supplementary Table 3.**
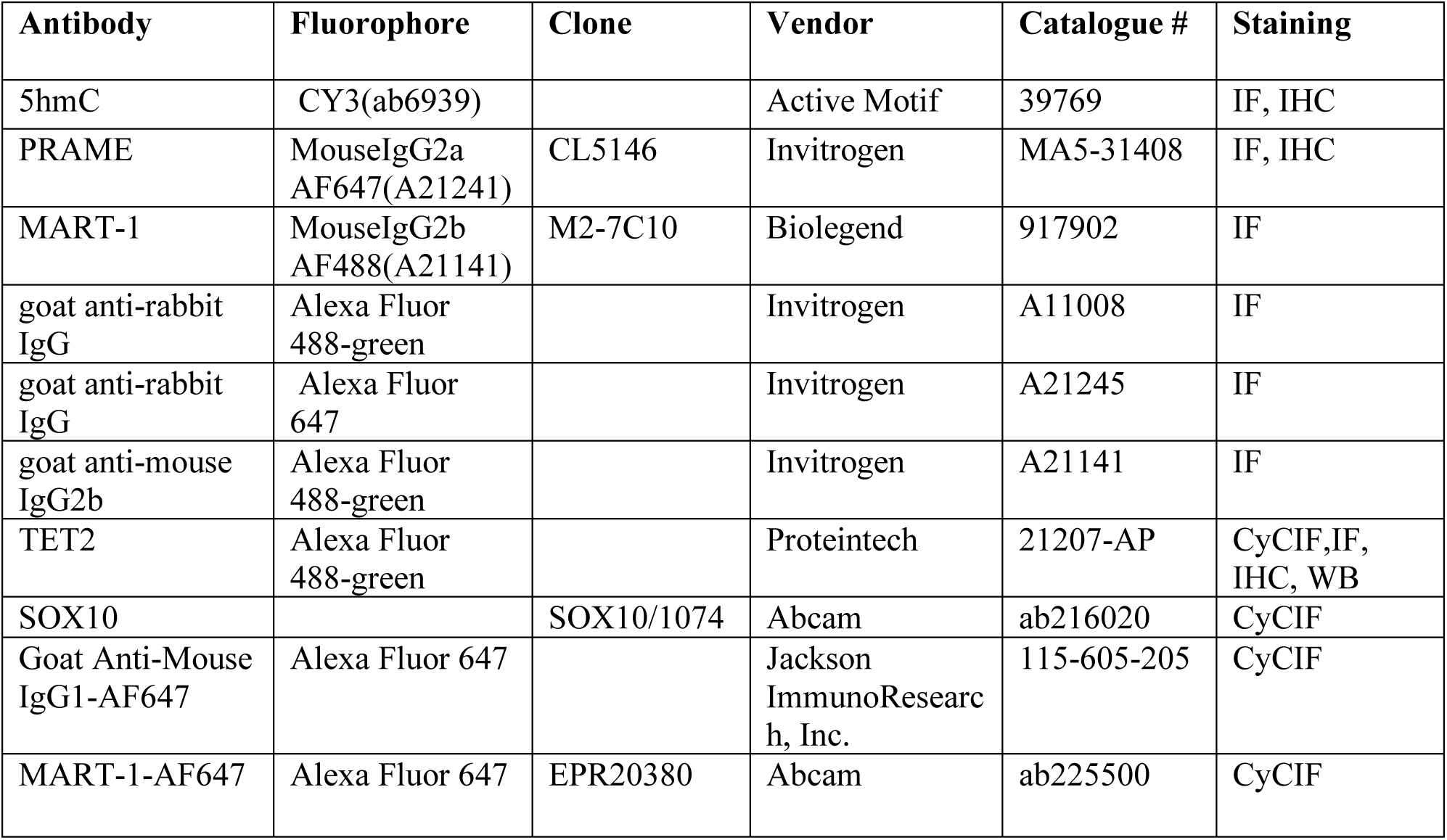
Antibody list of CyCIF, IF and IHC.

## Supplementary Figures

**SF1.**
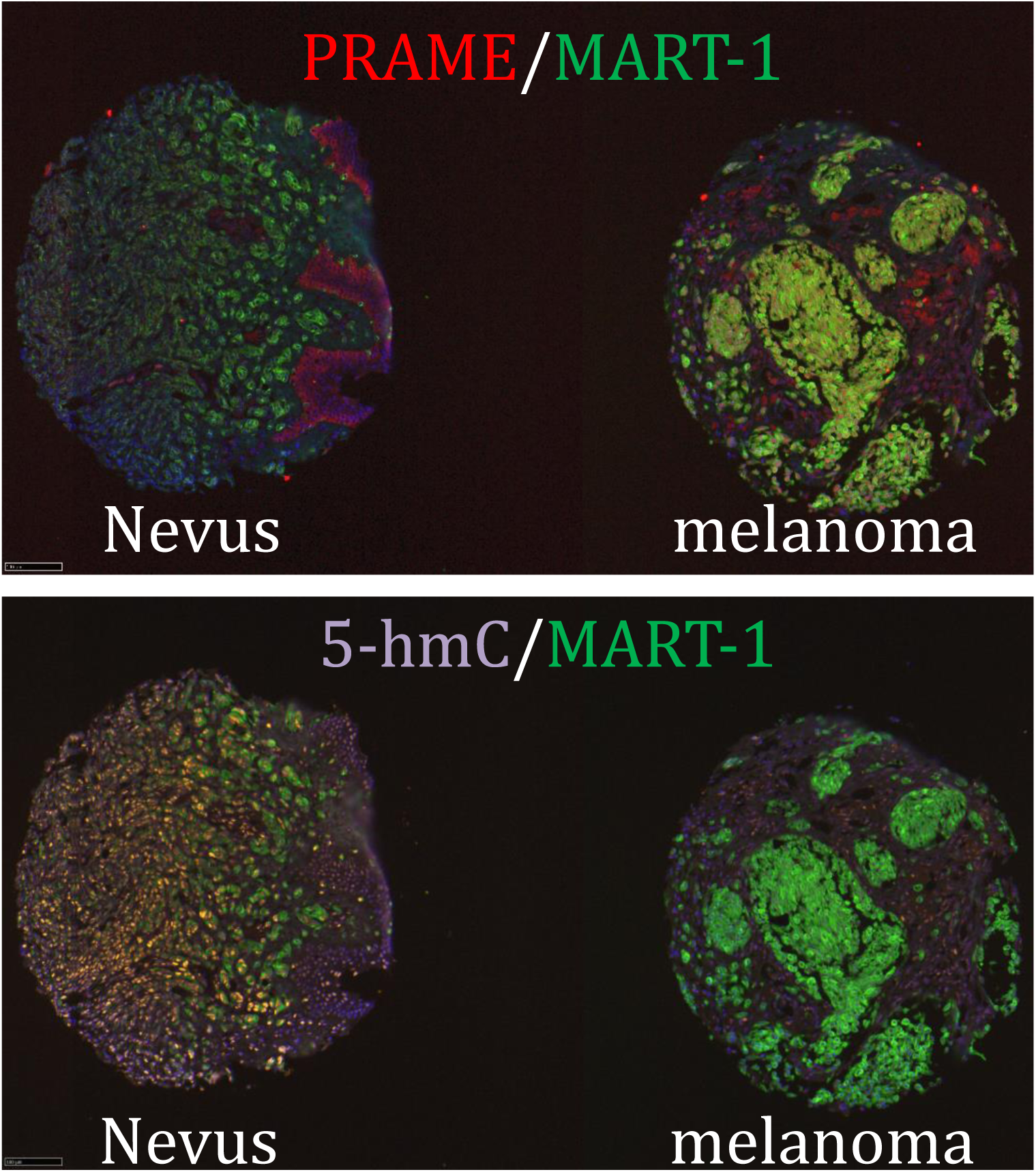
Multiplex immunofluorescence staining of nevus/melanoma tissue microarray.

**SF2.**
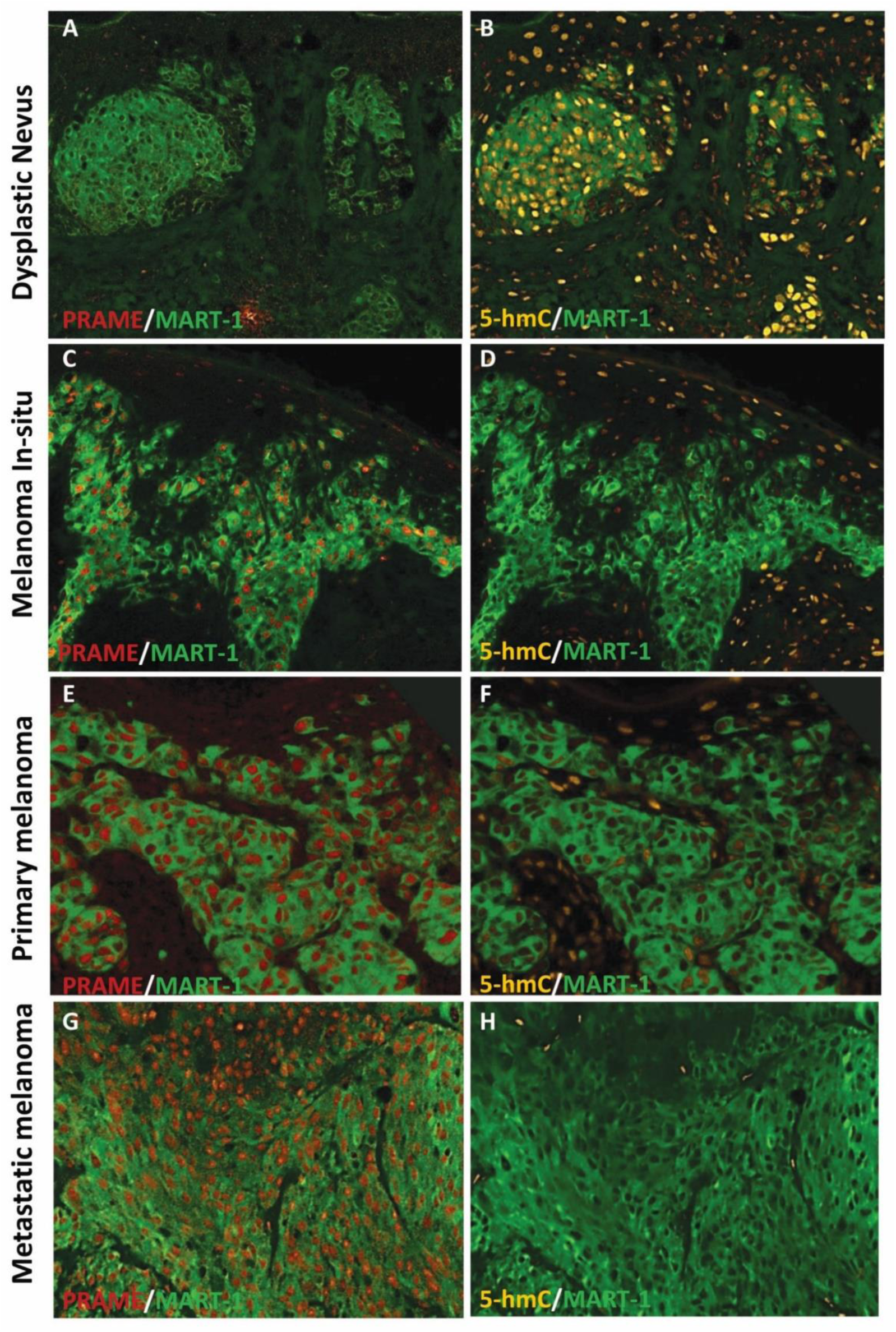
Multiplex immunofluorescence staining of nevus/melanoma tissue specimens.

## Acknowledgments

The findings published here are in part based from data obtained from the TCGA Research Network: https://www.cancer.gov.tcga

The Genotype-Tissue Expression (GTEx) Project was supported by the Common Fund of the Office of the Director of the National Institutes of Health, and by NCI, NHGRI, NHLBI, NIDA, NIMH, and NINDS.

## Ethics Approval and Consent to Participate

Ethics approval for this study was obtained from the Mass General Brigham Human Research Committee (IRB: 2015P001791) Informed consent was not necessary, as all tissue samples were deidentified. This study was performed in accordance with the Declaration of Helsinki.

## Author Contributions

CGL, GFM and PKS performed study concept and design; ED and SX provided acquisition of study samples. GF, SX, TV, RF, RP, IK, PS developed methodology, and performed analysis and interpretation of data, and statistical analysis. SX provided technical and material support. Authors AJZ, DVC, FA, RF and CGL wrote and reviewed the original manuscript draft. Substantial review and revision of the manuscript was performed by RF, TV, PKS, GFM and CGL. All authors read and approved the final paper.

## Funding Statement

This work was partially supported by NIH grants U2C-CA233262 (P.K. Sorger). A. J. Zhang receives a predoctoral stipend from an NIH T32 (T32TR004418).

## Data Availability Statement

All data needed to evaluate the conclusions in the paper are present in the paper and/or the Supplementary Materials. Additional data related to this paper may be requested from the authors.

## Outside activities

PKS is a co-founder and member of the BOD of Glencoe Software, and member of the SAB for RareCyte, NanoString, Reverb Therapeutics and Montai Health; he holds equity in Glencoe, Applied Biomath, and RareCyte. PKS consults for Merck and the Sorger lab has received research funding from Novartis and Merck in the past five years. The other authors declare no outside interests. GFM serves on the scientific advisory board of Biocoz Global and Stemson Therapeutics.

